# Learning performance and GABAergic pathway link to deformed wing virus in the mushroom bodies of naturally infected honey bees

**DOI:** 10.1101/2023.10.08.561384

**Authors:** Szymon Szymański, David Baracchi, Lauren Dingle, Alan S. Bowman, Fabio Manfredini

## Abstract

Viral infections can be detrimental to the foraging ability of the Western honey bee *Apis mellifera*. These include the deformed wing virus (DWV), which is the most common honey bee virus and has been proposed as a possible cause of learning and memory impairment. However, evidence for this phenomenon so far has come from artificially infected bees, while less is known about the implications of natural infections with the virus. Using the proboscis extension reflex (PER), we uncovered no significant association between a simple associative learning task and natural DWV loads. However, when assessed through a reversal associative learning assay, bees with higher DWV loads performed better in the reversal learning phase.

DWV is able to replicate in the honey bee mushroom bodies, where the GABAergic signalling pathway has an antagonistic effect on associative learning but is crucial for reversal learning. Hence, we assessed the pattern of expression of several GABA-related genes in bees with different learning responses. Intriguingly, mushroom body expression of selected genes was positively correlated with DWV load, but only for bees with good reversal learning performance. We hypothesize that DWV might improve olfactory learning performance by enhancing the GABAergic inhibition of responses to unrewarded stimuli, which is consistent with the behavioural patterns that we observed.

Our results suggest that previously reported DWV-driven learning deficits might be exclusive to acute, artificial infections and do not occur in naturally infected bees, stressing the importance of investigating more ecologically relevant scenarios when assessing host-parasite systems.

**Summary statement:** This study describes a virus-associated increase in learning in honey bees and proposes a mechanism based on GABA to explain the interplay between infection and cognition in the insect brain.

## Introduction

Honey bee colonies are susceptible to a wide range of parasites, and infections have been associated with incidences of colony death (Cox-Foster et al., 2007; Oldroyd, 2007; Cepero et al., 2014). While infections can lead to increased mortality, more subtle effects can also be detrimental for the colony. For example, infections with the microsporidian *Nosema ceranae* have been associated with less chances for bees to perform foraging duties or to carry pollen (Lach et al., 2015) and the Israeli acute paralysis virus has been shown to induce more frequent outbound flights from the hive by foragers in conjunction with a low-quality pollen provision (Dolezal et al., 2019). Also, it has been proposed that such infection-derived behavioural effects are crucial factors in over-wintering losses and colony death, as they affect the well-being and fitness of the entire colony (P. Chen et al., 2021; Decourtye et al., 2003).

Deformed wing virus (DWV) is the most common honey bee virus: infections can be lethal to individual bees as well as to the whole colony (Martin et al., 2015). DWV is strongly associated with the parasitic mite *Varroa destructor*, which acts as an efficient vector of the virus (Martin and Brettell, 2019; Chapman et al., 2023). Severe (overt) infections, usually acquired at the larval stage, cause a characteristic wing deformity in newly emerged bees (Martin and Brettell, 2019), alterations of the cuticular profiles (Baracchi et al., 2012), and can also cause additional symptoms such as bloated abdomens, ataxia, and discolouration (de Miranda and Genersch, 2010). However, more subtle consequences of DWV infections have also been reported, particularly when the virus is not transmitted by *Varroa* or persists in the population through chronic (covert) infections (Wells et al., 2016). Behavioural and cognitive impairments like deficits in olfactory learning and memory are among the reported symptoms associated with covert infections (P. Chen et al., 2021; Iqbal and Mueller, 2007; Tang et al., 2021). Although these effects of DWV are not lethal for the individual bee, they can be detrimental at the colony level, as they can impair the ability of workers to effectively forage, therefore threatening colony fitness and survival.

Several studies examining the effect of DWV on the physiology and cognition of honey bees have used artificial infections. In these works, the severity of achieved infections is not always reported and it transpires that different modes of infections (like feeding or injection) can produce contrasting results (P. Chen et al., 2021; Iqbal and Mueller, 2007; Tang et al., 2021). For example, Iqbal and Mueller (2007) reported virus-associated deficits in both learning and memory using DWV injections, while Chen et al. (2021) only reported deficits in long-term memory by feeding bees with DWV. The typical paradigm used in these studies to measure cognitive impairment is a change in the proboscis extension reflex (PER) response rate to conditioned odours. During PER, a bee will extend its proboscis in response to sugar being placed on its antennae. PER is readily associated with odours through classical conditioning: therefore, it is a widely used proxy for insect learning and memory (Giurfa and Sandoz, 2012). The system has also been broadly used to investigate reversal learning by conditioning bees to respond to two odours, and then reversing the reinforcements associated with them - sugar stimulation of the antennae or lack thereof (Mota and Giurfa, 2010). The link between performance in PER and the immune responses triggered by a parasitic infection has been well characterized (Mallon et al., 2003; Scheiner et al., 2003), including the molecular mechanisms underpinning the learning process (Raccuglia and Mueller, 2013). However, it remains unclear what the mechanisms are by which a virus like DWV can affect PER conditioning.

Some of the proposed mechanisms include alterations in olfactory perception, neuronal signalling, and carbohydrate metabolism pathways (P. Chen et al., 2021; Iqbal and Mueller, 2007; Pizzorno et al., 2021). One mechanism that has not been explored yet is GABAergic neuronal signalling within the mushroom bodies (MBs). The MBs are important integration centres of the insect brain with a key role in learning and memory (Homberg, 1984; Menzel, 2012), and GABA is known to play a role in modulation of these functions across many insect species (Dupuis et al., 2010). Experiments conducted in honey bees and *Drosophila melanogaster* fruit flies showed that the concentration of GABA in the MBs as well as the expression of GABA receptors and GABA synthesis genes have a strong antagonistic effect on insect associative learning (Liu et al., 2007; Liu and Davis, 2008; Raccuglia and Mueller, 2013). This is supported by the fact that in the calyces of the MBs, GABAergic feedback loops are crucial for reversal learning (Ganeshina and Menzel, 2001; Boitard et al., 2015). Intriguingly, DWV has been localised in the honey bee MBs – where it actively replicates (Shah et al., 2009) – and can induce changes in neuronal signalling (Pizzorno et al., 2021), suggesting a direct interaction between DWV and cognitive functions in this brain region that could be mediated by GABA.

Although GABA has not been assessed yet in response to DWV infections, this pathway has been proposed as a potential mechanism to explain the effects of pesticides on honey bee olfactory learning. Numerous types of pesticides are known to decrease the learning performance of bees in a dose-dependent and mode of exposure-dependent manner (Siviter et al., 2018; Carlesso et al. 2020; Cabirol et al., 2023). For example, early exposure to imidacloprid was associated with differential expression of GABA receptor subunit genes (Y. Chen et al., 2021), while exposure to thiamethoxam was associated with altered expression of synapsin, involved in the release of GABA in the central nervous system (Tavares et al., 2019). Moreover, nectar-borne GABA can directly affect learning and memory of honey bees (Carlesso et al., 2021) and bumble bees (Calderai et al., 2023).

In this study we used PER to test the associative and reversal learning abilities of honey bees with naturally occurring DWV infections. We then quantified DWV loads in the MB of bees that underwent PER, to investigate if cognitive impairments correlated with DWV loads. Finally, we assessed the expression of genes in the GABA pathway in MB of bees that showed different responses in reversal learning tests. Notably, our experiments used honey bees that were naturally infected with DWV, clearly separating our experimental approach from previous research where the virus was introduced *via* injection or oral administration. By using natural infections, we were able to test the effect of a virus on cognition and GABA-related gene expression in a more ecologically relevant context.

## Materials and Methods

### Colonies

Colonies of *A. mellifera* used in this study were kept in two locations near Aberdeen, Scotland: Cruickshank Botanical Garden, on the University of Aberdeen Kings College campus (Grid Reference NJ936085), and in Newburgh, Aberdeenshire (NJ998260). The Cruickshank colonies receive standard *Varroa* treatments and consequently have low *Varroa* infection rates and low DWV levels. The Newburgh colonies instead receive no *Varroa* treatments and have therefore relatively high Varroa and DWV levels (Woodford et al., 2022).

### Simple associative learning assay

Two colonies in Cruickshank Botanical Gardens and two in Newburgh were used as a source of bees for the simple associative learning trials. Foragers returning from foraging trips were captured between 09:00 – 10:00 at the colony entrance and transferred to the lab. Here, they were immobilised on ice for harnessing, fed with 3-5 μL of 30% (w/w) sucrose solution and maintained in the harness for an hour before the start of the assays. Harnessed bees were initially desensitised to water by touching the antennae with a soaked toothpick.

Bees were assessed with the Proboscis Extension Reflex (PER) absolute conditioning test. They were conditioned to respond to citral (Sigma-Aldrich, Saint Louis, Missouri, USA) through a 5-time exposure to the odour with reinforcement by touching the antennae with a 30% (w/w) sugar solution. 5 μL citral was pipetted onto a piece of filter paper, which was then inserted into a 20 mL syringe. The bees were exposed to citral by expressing the full volume of air from the syringe around 0.5 cm away from the bees’ antennae.

General PER olfactory conditioning protocol followed Mota and Giurfa (2010), with only one scent being used and no reversal performed. Foragers that exhibited PER to the first citral exposure were discarded. Foragers were then scored from 0 to 4 according to their responses during conditioning (0 = no response during all trials, 4 = PER response to 4 trials). Thirty minutes after the conditioning was concluded, foragers were exposed to the same odour as a mid-term memory test.

After behavioural testing, bees were frozen in a -80°C freezer and stored there until later processing for molecular work.

### Reversal associative learning assay

For this experiment, one hive in Cruickshank Botanical Garden and one in Newburgh were used – foragers were sampled, harnessed, and fed as described above. An hour after harnessing, bees were desensitised to water by touching the antennae with a soaked toothpick. Then, the bees were tested for sucrose responsiveness by exposure to six sucrose solutions of increasing concentration (0.1, 0.3, 1, 3, 10, 30% w/w) with an inter-stimulus interval (ISI) of 2 minutes (Carlesso et al., 2020). Sugar exposures were separated by water exposures to avoid sensitisation. To account for potential laterality in sucrose responsiveness, both antennae were stimulated (Baracchi et al., 2018). Individuals not responding to sucrose after exposure to a final 50% w/w solution and individuals responding to water after 30 min of desensitisation were excluded from the following steps. The sucrose responsiveness score (SRS) was calculated as the number of sucrose concentrations eliciting PER responses in bees.

After the initial tests, bees were fed 15 μL 30% (w/w) sucrose solution and maintained in the harness overnight. The following day, the differential conditioning experiment was conducted in two phases using two stimuli: odourant A (1-hexanol) and B (nonanal). The odour was delivered using a syringe, as described above. In the forward learning phase, A was reinforced positively, using 30% (w/w) sucrose solution placed on the antennae, and B was not reinforced (A = CS+, B = CS-; A+ and B-). In the reversal learning phase, the stimuli and reinforcements were switched, with A not reinforced and B positively reinforced with sucrose (A = CS-, B = CS+; A- and B+). In each phase, bees were exposed to the odorants five times per odour (total of 10 trials per phase) at an ISI of 10 minutes in a pseudorandomised sequence (i.e., at most two CS- or CS+ odours were presented in succession).

Specimens were divided into learners and non-learners according to the memory retention test. This test was performed 1 h after differential conditioning was concluded. Bees that responded with PER to odour B and did not respond to odour A were classified as learners since this was the last conditioning sequence they were exposed to (A-B+). All other bees were classified as non-learners.

After behavioural experiments were concluded, experimental bees were frozen in a -80°C freezer and stored there until later processing for molecular work.

### Analysis of data from learning assays

All statistical analyses were conducted in R, version 4.2.0 (R Core Team, 2017). Data were visualised using ggplot2 and ggpubr libraries. A standard significance threshold of 0.05 was adopted for all analyses.

PER scores in the simple associative learning assay were analysed using linear regression with the base R functions lm() and aov(). Tests were performed between PER score and log_10_-transformed mushroom body DWV loads, PER score and log_10_-transformed abdomen DWV loads, and also between mushroom body and abdomen log_10_-transformed DWV loads. Test assumptions were checked by examining residuals-fits plots and Anderson-Darling tests on residuals (ad.test() from nortest library). Differences in DWV loads according to performance in the mid-term memory test were assessed using a Wilcoxon rank test.

Bees’ learning performances in the reversal learning assay were analysed with a generalised linear mixed model (GLMM) (Baracchi et al., 2020). The GLMM was designed with a binomial error structure – logit-link function, glmer function of package lme4. The iterative algorithm BOBYQA was used to optimize the models. We retained the significant model with the highest explanatory power.

The selected model for the reversal learning phase included ‘bee response’ (1 = PER, 0= no PER) as the dependent variable, ‘CS’ and ‘group’ (i.e., learners or non-learners) as fixed factors, and ‘DWVlog’ and ‘conditioning trial’ as covariates. In all models, the individual identity ‘IDs‘ were entered as a random factor to account for repeated measures. The difference in DWV loads between learners and non-learners was assessed using a Mann-Whitney test.

### Quantification of DWV loads using RT-qPCR

We used a molecular assay to quantify viral loads in different tissues from honey bee samples collected after the Simple Associative Learning Assay (mushroom bodies and abdomens = 60 bees, abdomens alone = 126 bees) and Reversal Associative Learning Assays (mushroom bodies alone = 203 bees). Mushroom bodies were dissected on dry ice following Veiner et al. (2022) and homogenised in a Tissue Lyser device (QIAGEN) using 1 ml TRIzol and 2.3 mm Zirconia beads (Thistle Scientific, Glasgow, UK). Total RNA was isolated following routine TRIzol protocol and final elution was performed in 20 µl of nuclease-free water. Whole abdomens were processed in the same way, except for the final elution that was performed in 50 µl of water. To quantify viral loads, 300 ng of RNA was converted into cDNA using the iScript^TM^ cDNA Synthesis Kit (BioRad Laboratories Inc., Hercules, California, USA) and following the manufacturer protocol. cDNA samples were used as templates to run RT-qPCR reactions with three sets of primers: specific for strain A of the virus (DWV-A), specific for strain B (DWV-B) and global primers capable of detecting both strains (PAN primers). Primer sequences, RT-qPCR conditions, as well as viral load estimation methods were the same as described in Bradford et al. (2017). Throughout this work, DWV loads will be expressed in genome equivalents (GE).

Preliminary analyses revealed that there was a strong correlation between viral loads as detected with DWV-A and DWV-B primers and overall DWV loads as detected with PAN primers, both in the abdomen and mushroom body samples (Fig. S1, Table S2). Therefore, we opted to use only DWV loads as detected with PAN primers for this study.

### Sample selection for GABA gene expression assay

The learner group described above was subsampled to represent bees that had efficiently formed and reversed olfactory associations (Fig. 1): this subsample will be referred to as good reversers (GR) for simplicity. Similarly, the non-learner group was subsampled to obtain bees that efficiently formed olfactory associations but not reversed them (Fig. 1), which will be referred to as bad reversers (BR). The selection was made based on bees’ ability to pass or fail three behavioural criteria (details are shown in Table 1). This selection process resulted in a large number of samples being allocated to the BR group and only 17 samples to the GR group. Therefore, BR samples were pseudorandomly selected to also provide a matched sample size of 17.

**Figure 1.**
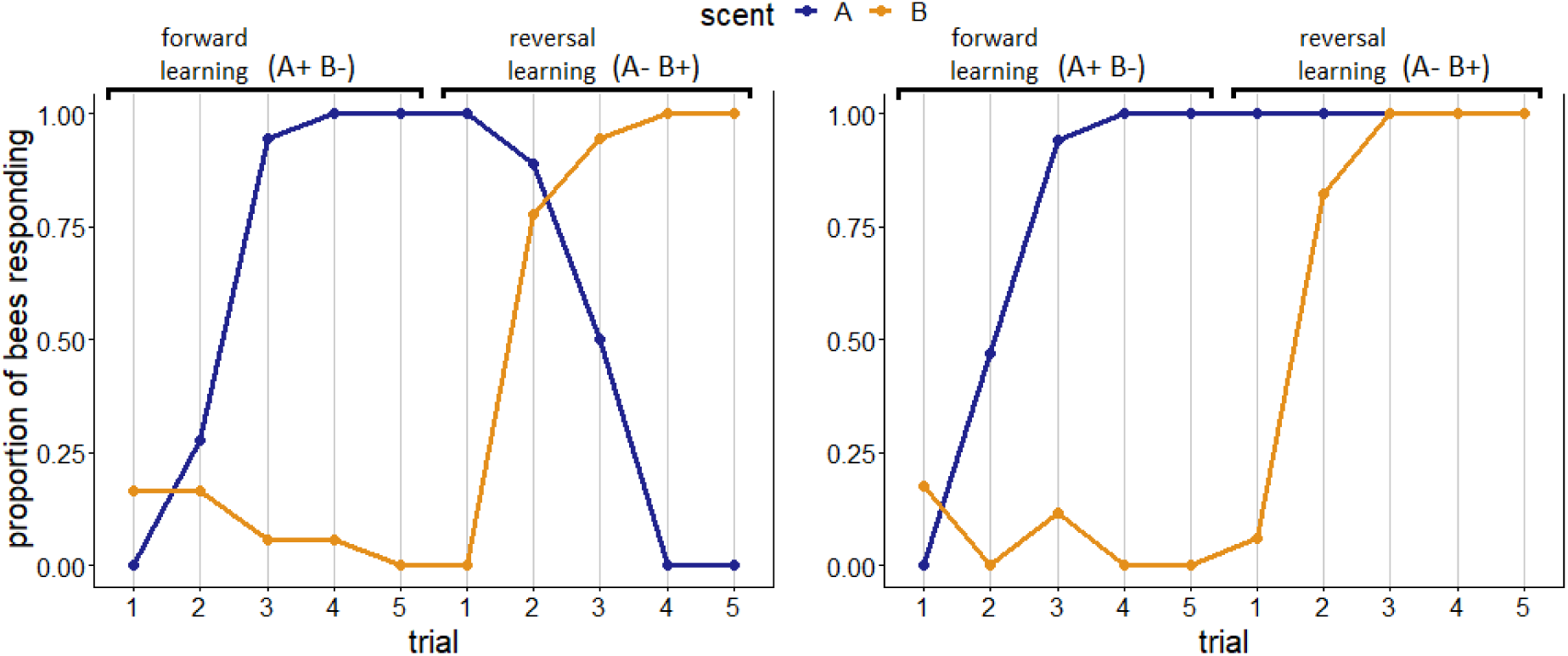
Performance of honey bees in a reversal associative learning assay. Responses graphs show the proportion of bees responding with PER to odours A (1-hexanol) and B (nonanal). The experiment was conducted in two phases, switching the reinforcement associated with odorants between the phases (A+ B- → A- B+). Left panel responses of bees belonging to the GR group (good learning ability and efficient reversal); Right panel: responses of the BR group (good learning ability, but not efficient reversal). N = 18 for both GR and BR groups. Subsets of bees represented in the figure were used for further gene expression analysis using RT-qPCR.

**Table 1.** Criteria used for subsampling learner and non-learner bees into good and bad reversers.

| Phase | GR (good reverser) | BR (bad reverser) |
| --- | --- | --- |
| *Forward learning* | PER for 5<sup>th</sup> trial of CS+ and no PER for 5<sup>th</sup> trial of CS- | PER for 5<sup>th</sup> trial of CS+ and no PER for 5<sup>th</sup> trial of CS- |
| *Reversal learning* | PER at most thrice during trials 2-4 for CS-.<br>No PER for 5<sup>th</sup> trial of CS- | --- |

### Preparation of RNA samples for gene expression analyses

Mushroom body RNA extracts were diluted with nuclease-free water up to 40 µL. Residual genomic DNA was digested using DNase-I (Zymo Research Corp, Irvine, California, USA) according to manufacturer protocol. After incubation at 23-25°C for 15 minutes, the RNA was purified and concentrated using the Zymo Research RNA Clean and Concentrator Kit following manufacturer protocol, with final RNA elution performed in 25 µL of nuclease-free water. Afterwards, samples were analysed with a NanoDrop^TM^ One Spectrophotometer (ThermoFisher Scientific, Waltham, Massachusetts, USA) to determine purity and yield.

To synthesise cDNA, aliquots constituting 250 ng of RNA were diluted with nuclease-free water to a volume of 15 µL and processed using the iScript^TM^ cDNA Synthesis Kit (BioRad Laboratories Inc., Hercules, California, USA) *as per* manufacturer protocol. Three additional RNA aliquots were processed without reverse transcriptase to provide non-enzyme controls (NECs).

### Expression analyses of honey bee GABA genes with RT-qPCR

The selected GABA-related genes (Table 2) were *Rdl* (ionotropic GABA-A receptor), *Camkii* (calmodulin-dependent protein kinase II), *Syn* (synapsin), *Abat* (4-aminobutyrate transferase) and *Gad1* (glutamate decarboxylase). Housekeeping genes used for normalisation were *RpS18* (ribosomal protein S18) and *RpL32* (ribosomal protein L32): these were sourced from Deng et al. (2020). A full description of the gene selection process as well as primer design and validation can be found in the supplementary materials. The complete candidate gene list is deposited in Table S1.

**Table 2.**
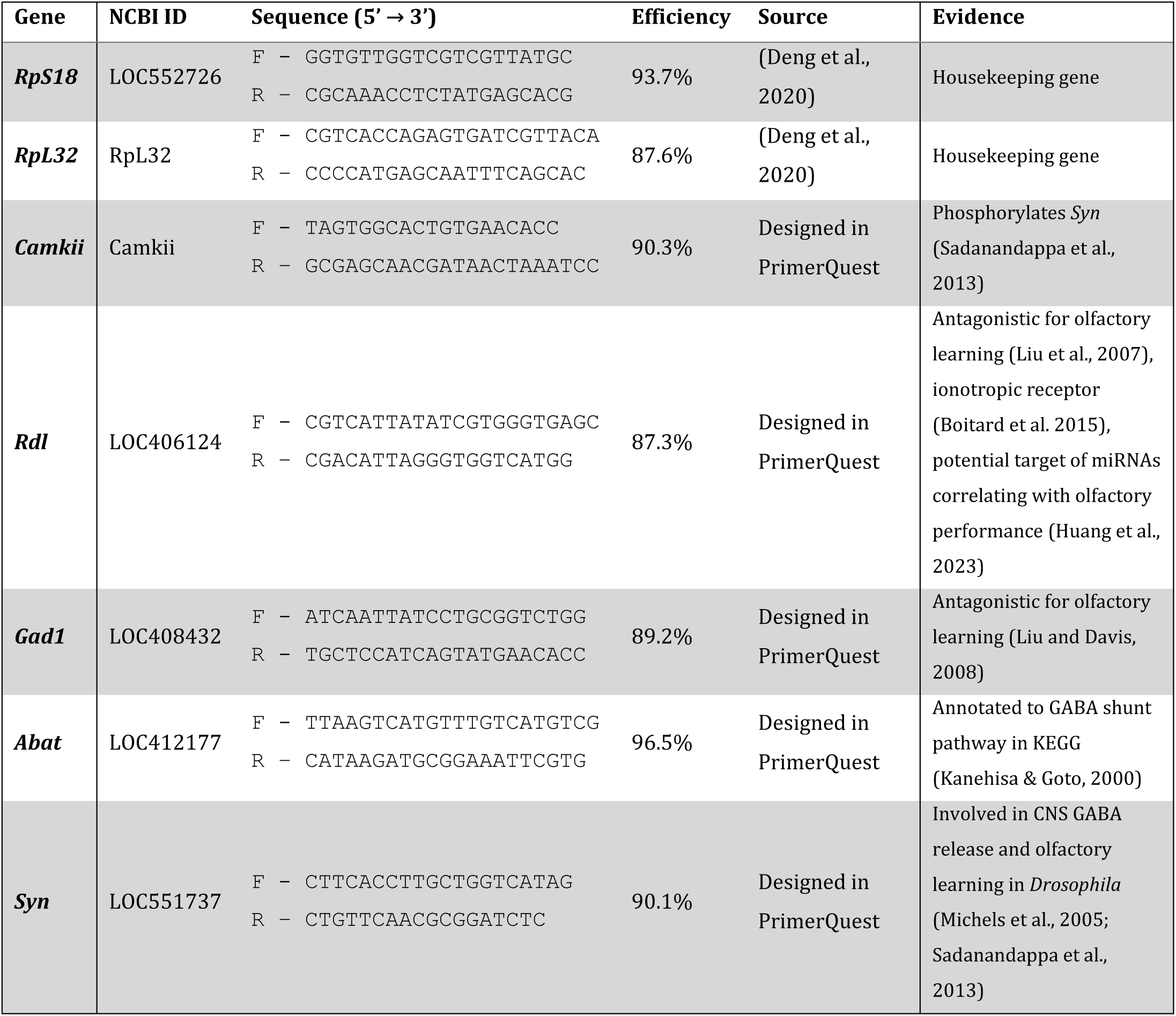
Gene list with primer pairs used for qPCR.

A total of 34 samples (GR and BR combined) were assayed in duplicates, with each gene tested on a separate 96-well plate for all samples, including 3 NECs and 3 water controls to check for non-specific amplification. Assays were performed on a BioRad C1000 Touch^TM^ Thermal Cycler with the CFX96^TM^ Optical Reaction Module attachment. Plates were pre-incubated at 95°C for 3 minutes, followed by 40 amplification cycles (95°C for 3s, 60°C for 20s, 72°C for 3s), finishing with a 4-minute elongation at 72°C. Data from the thermocycler were imported into BioRad CFX Manager software to examine melting curves and assess specificity of amplicons; thereafter, C_q_ values were exported for further data processing and statistical analyses with R. Sample duplicates were averaged and the C_q_ data were normalised using a modified Pfaffl method, which allows for the use of two reference genes (Hellemans et al., 2007; Vandesompele et al., 2002).

Prior to the analysis, all data for one sample in the BR group and one in the GR group were removed due to extreme expression values across all genes. For *Gad1* data, an additional GR sample had to be removed due to a lack of replication in the RT-qPCR assay. Differences in GABA-related gene expression between GR and BR groups were tested using separate 2-sample T-tests for each gene, as the assumptions for nested ANOVA could not be met. This was achieved using the base function t.test(). Repeated testing was accounted for using a Bonferroni correction (VanderWeele and Mathur, 2019). The assessment of T-test assumptions was performed using ad.test() and levene_test() functions from nortest and rstatix libraries respectively.

Changes in gene expression according to DWV loads were assessed using a generalised linear model with the R function glm(). The explanatory variables were ‘MB log_10_ DWV count’, ‘behavioural group’ (GR or BR), ‘gene’, as well as interaction terms between ‘DWVlog10*behavioural group’, and ‘DWVlog10*gene’. A Gamma log link function was used for this analysis because expression data are continuous and non-negative. Validity of the model was checked with an analysis of deviance using the base function pchisq(), by subtracting the model deviance from null deviance and testing the value against a χ^2^ distribution for the model’s degrees of freedom. The model was able to explain a significant proportion of the data deviance (χ^2^ = 23.15, df = 11, p = 0.017). A null model was also a worse fit for the data as demonstrated by the AIC value (482.62 for the model presented; 491.60 for the null model).

## Results

### Performance in simple learning does not correlate with viral loads

Linear regression analyses showed that DWV loads of neither mushroom bodies nor abdomens explained a significant proportion of variation in PER score data derived from simple associative learning (Table 3). Analogously, the performance of bees in the mid-term memory test was not significantly associated with DWV loads (Wilcoxon; W = 226, p-value = 0.648). However, mushroom body loads were significantly associated with abdomen DWV loads (Fig. 2). Both the slope and the intercept of this fit were significantly different from 0 (one sample t-test; slope T = 6.28, p = 4.73⨯10^-8^; intercept T = 7.67, p = 2.17⨯10^-10^). Regression assumptions were met, with small departures from normality, which were deemed not detrimental to the analysis.

**Figure 2.**
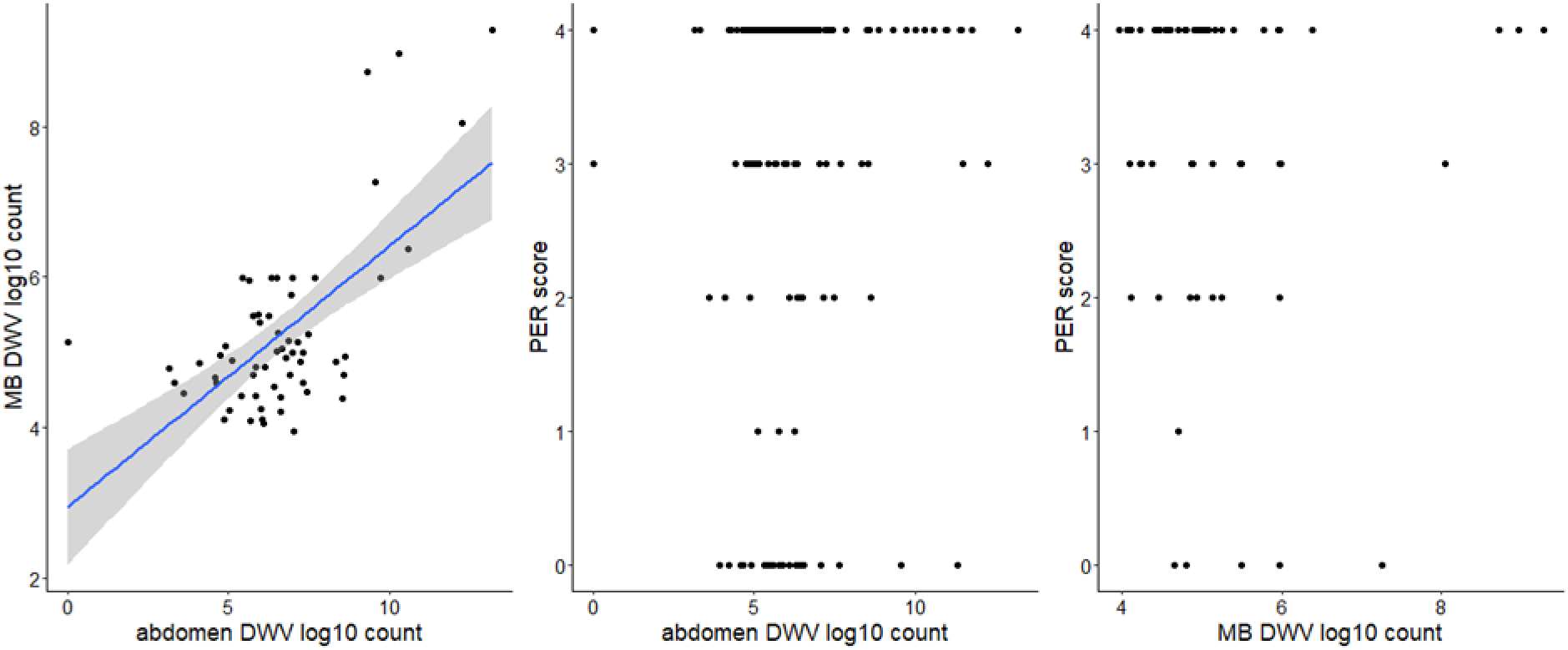
Regression analyses between PER score and DWV load data in a simple associative learning test. Scatterplots of mushroom body DWV counts against abdomen DWV counts (left), PER score against mushroom body DWV counts (centre) and PER score against abdomen DWV counts (right). A total of 186 bees were assessed using the associative learning assay. Neither abdomen nor MB DWV loads were significantly associated with performance (PER score) in this assay. Only mushroom body and abdomen DWV counts were significantly associated according to linear regression analysis (F_1,58_ = 39,41, p = 4.73⨯10^-8^). Line of fit is y = 2.985 + 0.346x. n = 60 for left and right panels; n = 186 for centre panel.

**Table 3.** Results of linear regression analyses of PER score, and mushroom body / abdomen DWV loads.

| Response variable | Explanatory variable | F-value | p-value | Df | sig | slope | intercept |
| --- | --- | --- | --- | --- | --- | --- | --- |
| PER score | abdomen DWV log<sub>10</sub> count | 0.31 | 0.576 | 1, 184 | ns | - | - |
| PER score | mushroom body DWV log<sub>10</sub> count | 0.04 | 0.840 | 1, 58 | ns | - | - |
| mushroom body DWV log<sub>10</sub> count | abdomen DWV log<sub>10</sub> count | 39.41 | 4.73 × 10<sup>-8</sup> | 1, 58 | *** | 0.346 | 2.985 |

### Performance in reversal learning positively correlates with viral loads

In the forward learning phase of the reversal associative learning assay (A+ B-), bees increased their PER response to the odours throughout the conditioning trials (GLMM, *sequence*; χ^2^ = 26.34, df = 1, p = 2.86⨯10^-7^) and, at the same time, discriminated the CS+ and CS-stimuli (GLMM, *CS*; χ^2^ = 23.70, df = 1, p = 1.12⨯10^-6^; Fig. 3C). DWV counts did not affect forward learning (GLMM, *DWVlog;* χ^2^ = 1.43, df = 1, p = 0.23; Figure 3A).

**Figure 3.**
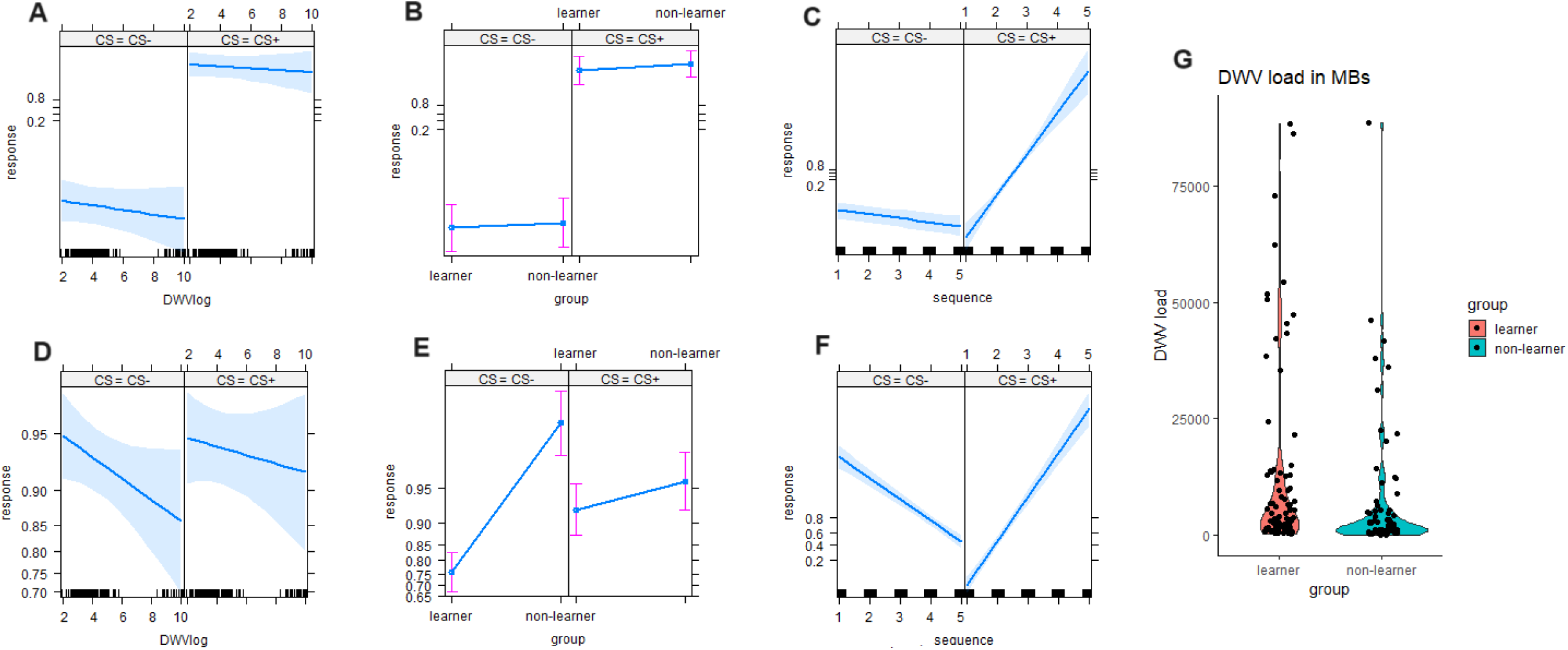
Correlation between reversal learning and DWV loads in honey bee mushroom bodies. Effect plots for the GLMM used to analyse behavioural responses during reversal learning assay (A-F) and a violin plot comparing DWV loads between behavioural groups (G). A total of 227 bees were processed in this assay (n = 114 learners; n = 89 non-learners). Panels A-C present effects during the forward learning phase (odour A = CS+, odour B = CS+) of the experiment and panels D-E present effects during the reversal phase (odour A = CS-, odour B = CS+). A: Effect of DWV load on responses to odours A (CS+) and B (CS-). B: Effect of behavioural group on response to odours A (CS+) and B (CS-). C: Effect of sequence (trial number) on responses to odours A (CS+) and B (CS-). D: Effect of DWV load on responses to odours A (CS-) and B (CS+). E: Effect of behavioural group on response to odours A (CS-) and B (CS+). F: Effect of sequence (trial number) on responses to odours A (CS-) and B (CS+). G: Differences in DWV loads between bees that passed the memory test (learners) and bees that did not pass the memory test (non-learners); the difference was significant (W = 4592, p-value = 0.224).

In the reversal learning phase of the experiment (A- B+), bees updated their information: they increased their PER response to B+ and decreased the response to A- (GLMM, sequence*CS; χ2 = 293.43, df = 1, p = 2.2⨯10-16; Fig. 3F). However, while the response to stimulus B+ was not influenced by DWV counts (Tukey, post-hoc test, p = 0.17), the response to stimulus A- was negatively associated with increasing DWV counts (GLMM, *DWVlog*CS*; χ2 = 12.67, df = 1, p = 0.0004; post-hoc test p < 0.0001; Fig. 3D). Similarly, learners and non-learners responded equally well to B+ but not to A-, as non-learners failed to stop responding to the unrewarded stimulus over the reversal training (GLMM, Group*CS; χ2 = 47.36, df = 1; p = 5.88⨯10-12; Fig. 3E).

By contrast, DWV loads differed between learners and non-leaners (Mann-Whittney; W = 1078, p = 0.007; Fig. 3G). Specifically, learners had higher DWV counts in their mushroom bodies than non-learners. SRS did not differ between learners and non-learners (Mann-Whitney U test; W = 4592, p-value = 0.224). Also, in this case, SRS did not correlate with DWV loads (Spearman test; r=-0.06, p-value = 0.41).

### Expression of GABA-related genes does not differ between behavioural groups

All groups of data were normally distributed, except for *Abat* in the GR group (Anderson-Darling test; A = 0.975, p = 0.010; all other tests p > 0.15). Equality of variances was also satisfied (Levene’s test, all tests p > 0.80). Therefore, T-tests were deemed a valid analysis and a small departure from normality should not be detrimental to it.

None of the intra-gene group comparisons proved significant (Table 4) Thus, the expression of GABA-genes in the mushroom body does not differ between good and bad reversers. Overall, data for *Abat* had lower dispersal than the other genes presented (Fig. 4). A common pattern across all genes was a higher expression in the BR group, however, this trend was not significant.

**Figure 4.**
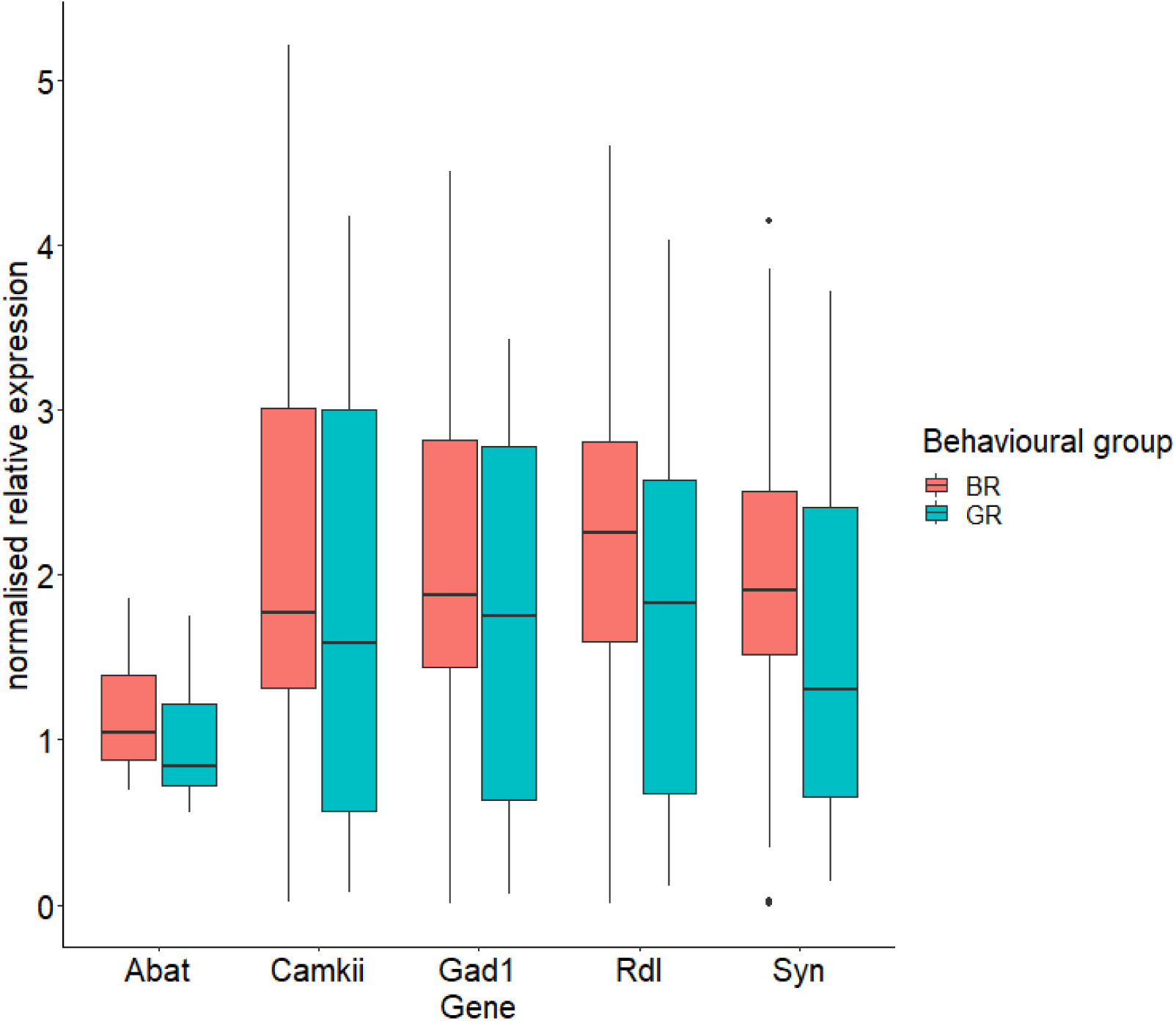
Relative expression of GABA-related genes in the mushroom bodies of bees exposed to reversal learning. Boxplots show median and quartile boundaries. Gene expression was assessed for a total of 34 bees subjected to a reversal learning assay. No significant differences were detected between behavioural groups for any gene (two-sample, two-tailed T-test); Bonferroni adjustment was used to account for multiple comparisons. n = 16 BR for all genes; n = 16 GR for *Abat, Camkii, Rdl, Syn*; n = 15 GR for *Gad1*.

**Table 4.**
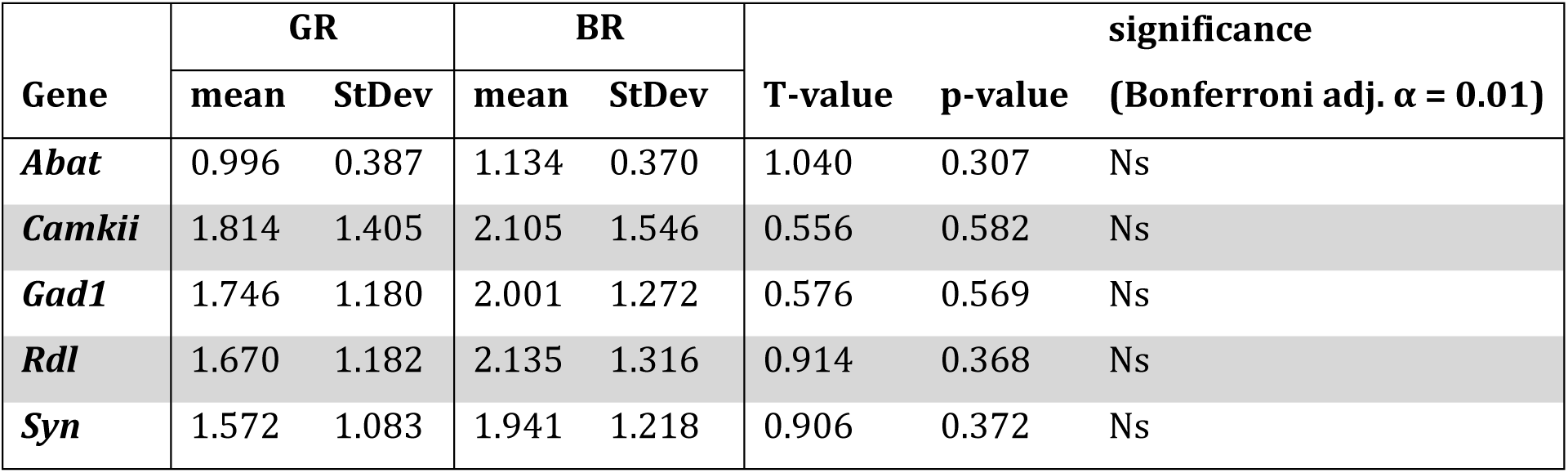
Descriptive statistics for normalised gene expression, showing results of two-sample t-tests between behavioural groups.

### Expression of GABA-related genes positively correlates with viral loads

The effect of DWV load on GABA-related gene expression was highly significant in GR bees (GLM, *DWVlog10*:*GR*; β est. = 0.473, StErr = 0.108; T = 4.365, p = 2.28⨯10^-5^) while this was not the case for BR bees (GLM, *DWVlog10*; β est. = -0.248, StErr = 0.134; T = -1.849, p = 0.067). Overall, among GR bees, the expression of all GABA-related genes seemed to increase with increasing log_10_ DWV load (Fig. 5). In our statistical model the gene *Abat* was used as a baseline and the effect of DWV load on expression was significantly different only for *Rdl* as compared to that baseline (GLM, *DWVlog10*:*Rdl*; β est. = 0.343, StErr = 0.170; T = 2.017, p = 0.046). Nevertheless, when the other genes were compared to *Abat* they achieved similar β coefficient estimates and were close to significance (detailed coefficients are shown in Table S3).

**Figure 5.**
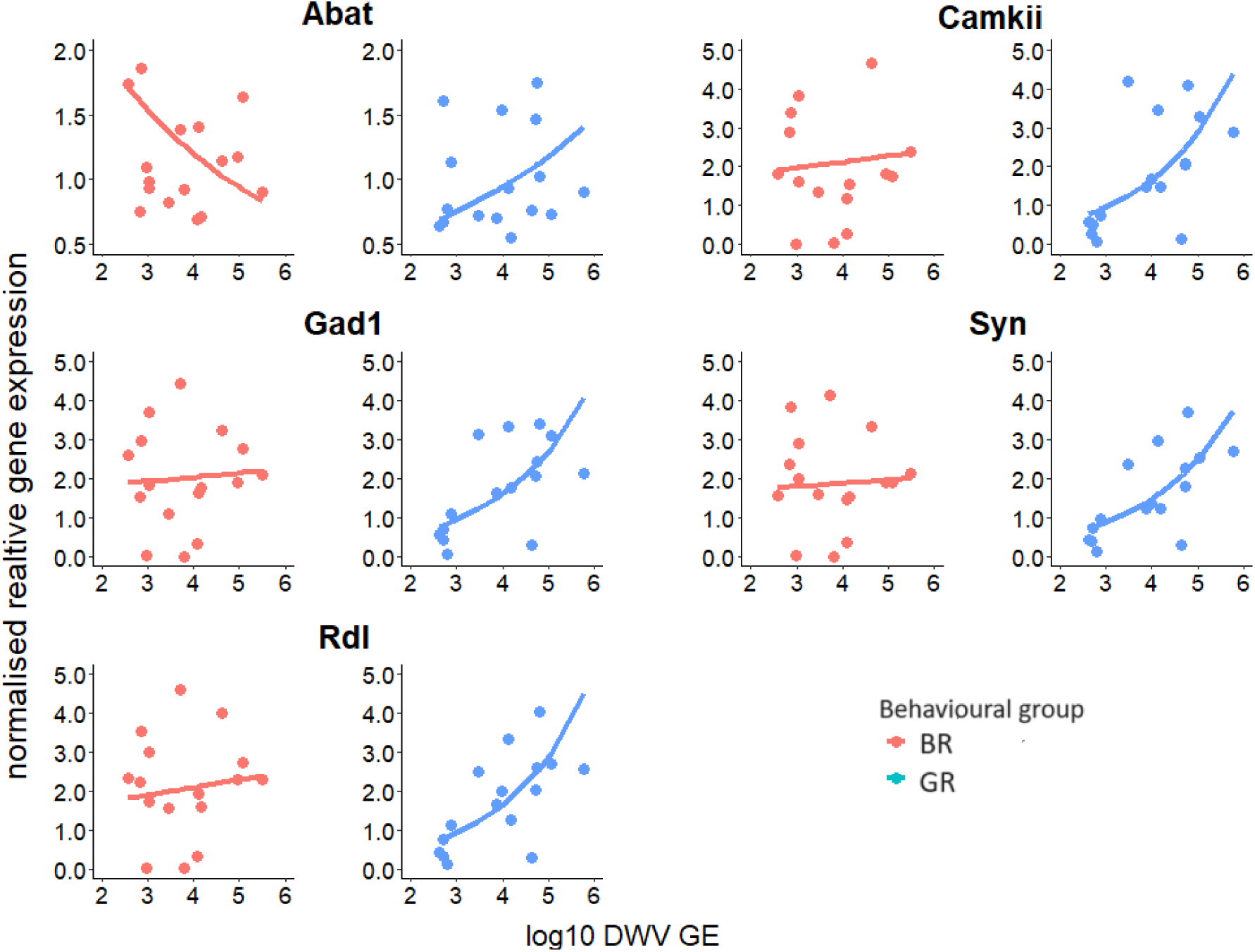
Correlation between GABA gene expression and DWV loads in honey bee mushroom bodies. Scatterplots show normalised relative expression of five genes related to GABA in the mushroom bodies of bee foragers (y-axes), and their mushroom body DWV loads (x-axes). Lines show fitted values produced by a multivariate, log-linked generalised linear model. Significant differences were found in slopes of fitted values (β est. = 0.473, StErr = 0.108; T = 4.365, p = 2.28⨯10^-5^) between GR and BR data. n = 16 BR for all genes; n = 16 GR for *Abat, Camkii, Rdl, Syn*; n = 15 GR for *Gad1*. Please note that *Abat* is on a different y-scale for visualisation purposes.

## Discussion

Here we combined a set of simple and reversal associative learning tests to characterize the cognitive performance of honey bee foragers naturally infected with DWV and spanning a broad range of viral loads. We then investigated the expression levels of a selected pool of GABA-related genes to assess the effect of chronic infections with DWV on these key regulators of neural functions. The performance of bees in the simple associative learning test did not significantly vary in response to DWV: this held true for viral loads in both mushroom bodies and abdomens. Mushroom body DWV loads were significantly predicted by abdomen DWV loads, which deems our use of mushroom body loads throughout this work representative of overall loads. In the reversal learning assay instead, learners had significantly higher DWV than non-learners. In an analysis of behavioural responses, bees with a higher DWV had lower responsiveness to CS- in the reversal phase, thus enhancing their learning performance by inhibiting the response to an unrewarded stimulus. Good and bad reversers did not differ in expression levels for the tested GABA-related genes. However, the expression of those genes increased with DWV load in good reversers.

The fact that PER learning score was not associated with either mushroom body or abdomen DWV loads can be explained as follows: DWV does not impair simple associative learning. Such a finding is in direct opposition to previous work, where DWV infection induced learning deficits in a similar assay (Iqbal and Mueller, 2007). This discrepancy may be explained by some key differences in the methods that the two studies adopted, as Iqbal and Mueller (2007) artificially infected adult bees using injections of DWV, while our study assessed bees that were naturally infected (very likely since larval or pupal stage). This is a key difference from several points of view: the modality of transmission of a viral parasite (i.e., artificial injection vs. natural infection) can dictate how the virus *behaves* in the host (e.g., viral replication) and can also affect how the host responds (Gisder et al., 2018), as piercing the cuticle with a syringe to deliver the lysate very likely triggers a different chain of reactions than acquiring the virus naturally through parasitisation by *Varroa* or contact with infected nestmates. In terms of the viable viral loads in the two studies, a comparison is impossible as Iqbal and Mueller (2007) did not report viral loads in infected bees or DWV lysate used for their experiments. Interestingly, experiments using artificial DWV infections such as Iqbal and Mueller (2007) can achieve markedly higher DWV loads than natural infections. For example, one study reported mean DWV log_10_ genome equivalents = 10 in artificially infected bees (Pizzorno et al., 2021), which is on par with the most severe natural infections observed in our bee colonies. Although Pizzorno et al. (2021) quantified their viral loads differently than us, it seems plausible that infection levels in Iqbal and Mueller (2007) were overall significantly higher than observed in our study, and a DWV-driven learning deficit might be specific to such extremely severe, acute infections. This is partially supported by another study (P. Chen et al. 2021) where bees were fed DWV lysate and infection levels in the brain reached 10^5^-10^6^ DWV copy numbers at 100 hours post-infection (thus comparable to our study). There, the conditioning responses of the control and infected bees were strikingly similar, and they only differed in long-term memory (P. Chen et al., 2021). Furthermore, it has already been demonstrated that some symptoms of DWV infection, such as wing asymmetry, develop in a viral load-dependent manner (Brettel et al., 2017), lending additional support to this hypothesis.

In contrast with the patterns observed for simple associative learning in our assay, individuals with higher levels of infection performed better in the reversal learning assay. This effect was primarily a result of their increased inhibition of responses to previously rewarding stimuli (i.e., bees with higher DWV showed lower responsiveness to negative stimuli). Such a result also explains why no effect on performance was seen in the simple associative learning assay: simply, there was no negative stimulus in the associative learning assay, as one odour was used, and it was rewarded throughout the assay. Thus, if DWV-associated learning enhancement is driven by inhibition of responses, the effect simply could not be observed in the associative learning assay.

Expression of GABA-related genes did not differ between good and bad reversers. This is in opposition with what we initially hypothesised, as previous research has shown that activation of ionotropic GABA receptors is necessary for successful reversal learning (Boitard et al., 2015). Indeed, the experimental inhibition of mushroom body GABAergic signalling was shown to completely stop reversal learning in honey bees (Devaud et al., 2007). Our selection of candidate genes includes not only ionotropic receptors (Rdl) but also genes involved in the synthesis (Liu and Davis, 2008) and release of GABA (Liu et al., 2007), and therefore these results do not seem to support any link between expression of GABA-related genes and an individual’s ability or inability to perform reversal learning. One aspect worth of future investigation will be to test if the facilitation of reversal learning by GABA responds to a threshold effect, so that a minimum degree of GABAergic signalling is necessary for reversal learning, and once the degree is reached learning continues to happen for a certain period. In this scenario, the levels of expression of GABA-related genes at the specific time point that we assessed might not reflect the overall activation status of GABAergic pathways, and a more comprehensive time series of observations would be required to explore whether there is any support for this hypothesis.

It is interesting to note that, although not statistically significant, the bad reverser group consistently showed higher expression of all tested GABA-related genes. One explanation for this pattern might be that gene activation reflects the fact that bad reverser bees had not completed the associative learning process when sampled, as their responses were not consistent with the stimuli presented at the end of the assay. On the contrary, good reverser bees could have successfully learned the association and were retrieving it from their memory – hence activation of GABA-related genes was no longer necessary. GABA is known to modulate learning in the mushroom body in an intricate time-dependent manner, and in a PER paradigm it is only effectively antagonistic of learning during the few seconds of odour presentation (Raccuglia and Mueller, 2013). Notably, GABA in the mushroom bodies has been demonstrated to not modulate memory retrieval at all (Raccuglia and Mueller, 2013). Therefore, in such a scenario, slightly higher GABA expression in bad reverser bees could be justified with these bees not having completed the learning process yet, and good reversers bees having retrieved the memorised association by the time their memory was tested.

A key result of our study was the positive correlation between expression of the selected GABA-related genes and DWV loads in the mushroom bodies. This was observed for all investigated genes but, intriguingly, it held true only for good reverser bees. Potentially, such an increased GABA-gene expression could allow for a more efficient inhibition of responses to a previously rewarded stimulus. This is supported by the behavioural results of this assay, as learner bees (good reversers are a subsample of these) expressed a negative association between DWV loads and responsiveness to CS- in the reversal phase of the reversal associative learning assay. Furthermore, such an interpretation is congruent with the biological functions of GABA in the mushroom body during the learning process. As outlined above, it is known that mushroom body GABA overwhelmingly inhibits learning in insects. It was experimentally proven that an increased concentration of GABA, or expression of GABA receptors and synthesis genes, are all able to stop the learning processes within the mushroom bodies (Liu et al., 2007; Liu and Davis, 2008; Raccuglia and Mueller, 2013). It may be assumed that the GABAergic pathway also inhibits responses to previously learned stimuli once they stop being rewarded. Thus, it is possible that DWV acts to enhance reversal learning performance through the stimulation of GABA pathways in the mushroom bodies.

Our study represents a first step in the understanding of the complex interaction between viral infections, learning and gene expression in the honey bee mushroom bodies. At the same time, the results presented here highlight some possible limitations to this work. First, whole mushroom body samples might not provide the appropriate anatomical resolution to demonstrate GABAergic differences in gene expression between bees with different reversal learning abilities. The anterior paired lateral neuron is a GABAergic neuron that projects from the antennal lobe into the mushroom body. This neuron is known to form a memory trace during associative learning and the manipulation of *Gad1* (GABA synthesis enzyme) expression in the neuron modulates learning ability in *Drosophila* (Liu and Davis, 2008). Expression patterns of *Rdl* (ionotropic GABA-A receptor) in the mushroom body closely follow the projections of that neuron (Liu et al., 2007). Therefore, the GABA-ergic mediation of olfactory reversal learning may occur at the level of projection neurons and not directly within the mushroom body. Future works could thus utilise methods with a higher resolution, such as single-cell sequencing, to distinguish between different neuron types in and around the mushroom body (Zhang et al., 2022). Another limitation, already highlighted in this discussion, pertains to the focus on a single time point for screening GABA-related gene expression. Evidence shows that GABA might operate in a highly time-dependent fashion (Raccuglia and Mueller, 2013), and therefore a wider range of sampling time points should be tested in the future to provide a more comprehensive characterization of the expression patterns of GABA-related genes. Finally, a wider range of natural DWV infections should be incorporated when investigating the effect of viruses on honey bee learning. Our study represents a first step in this direction, highlighting how the effect of natural infections on honey bee cognitive abilities might present significant differences from what previously reported for artificial infections. This testifies to the importance of expanding the assessment to a range of modalities of infections, which will provide new insights into this fascinating host-parasite system and a better understanding of its relevance in an ecological context.

## Abbreviations

BR: bad reverser
DWV: deformed wing virus
GE: genome equivalent
GR: good reverser
MB: mushroom body
PER: proboscis extension reflex
SRS: sucrose responsiveness score

## Acknowledgements

The authors extend their thanks to Dr Craig Christie for unlimited support with laboratory techniques, to Dr Ewan Campbell for beekeeping support and to Estefania Hugo Arnábal for help in primer design.

## Competing interests

We declare no competing interests.

## Author contributions

Conceptualisation: A.S.B., D.B., F.M., S.S.

Methodology: D.B., F.M., L.D., S.S.

Investigation: L.D., S.S.

Resources: A.S.B., F.M.

Formal analysis: D.B., S.S.

Visualization: D.B., S.S.

Writing – original draft: S.S.

Writing – review & editing: A.S.B., D.B., F.M., L.D., S.S.

Supervision: A.S.B., F.M.

Funding acquisition: A.B., F.M.

## Funding

This work was funded by a Research Grant awarded by the C. B. Dennis British Beekeepers’ Research Trust to F.M. and A.S.B: this grant also contributed salary to L.D. for the whole duration of the project. Additionally, D.B. acknowledges the support of NBFC to the University of Florence, Department of Biology, funded by the Italian Ministry of University and Research, PNRR, Missione 4 Componente 2, “Dalla ricerca all’impresa”, Investimento 1.4, Project CN00000033.

## Data availability

All relevant data can be found in the main text or in the supplementary materials. No custom software has been used in this work. Preliminary checks of qPCR data were conducted in BioRad CFX Manager^TM^ version 3.0, which is available from the manufacturer. All data analyses were conducted in RStudio, an open-source interface, using R (version 4.2.0) packages and libraries available from public repositories.

## Notes

### Competing Interest Statement

The authors have declared no competing interest.

## References

Baracchi, D., Cabirol, A., Devaud, J. M., Haase, A., d’Ettorre, P. and Giurfa, M. (2020). Pheromone components affect motivation and induce persistent modulation of associative learning and memory in honey bees. Commun. Biol. 2020 3:1, 3(1), 1–9.

Baracchi, D., Fadda, A. and Turillazzi, S. (2012). Evidence for antiseptic behaviour towards sick adult bees in honey bee colonies. J. Insect Physiol., 58(12), 1589–1596.

Boitard, C., Devaud, J.-M., Isabel, G. and Giurfa, M. (2015). GABAergic feedback signaling into the calyces of the mushroom bodies enables olfactory reversal learning in honey bees. Front. Behav. Neurosci., 9(JULY), 198.

Bradford, E. L., Christie, C. R., Campbell, E. M. and Bowman, A. S. (2017). A real-time PCR method for quantification of the total and major variant strains of the deformed wing virus. PLoS One, 12(12), e0190017.

Brettell, L. E.. Mordecai, G. J., Schroeder, D. C., Jones, I. M., Da Silva, J. R., Vicente-Rubiano, M., and Martin, S.J. (2017). A Comparison of Deformed Wing Virus in Deformed and Asymptomatic Honey Bees. Insects 2017, 8, 28.

Cabirol, A., Gómez-Moracho, T., Monchanin, C., Pasquaretta, C. and Lihoreau, M. (2023). Considering variation in bee responses to stressors can reveal potential for resilience. J. Appl. Ecol. 60, 1435–1445.

Calderai, G., Baggiani, B., Bianchi, S., Pierucci, V., Foti, M., Scibetta, F. and Baracchi, D. (2023). Nectar-borne GABA promotes flower fidelity in bumble bees. Entomol. Gen.

Carlesso, D., Smargiassi, S., Pasquini, E., Bertelli, G. and Baracchi, D. (2021). Nectar non-protein amino acids (NPAAs) do not change nectar palatability but enhance learning and memory in honey bees. Sci. Rep. 2021 11:1, 11(1), 1–14.

Carlesso, D., Smargiassi, S., Sassoli, L., Sassoli, L., Cappa, F., Cervo, R. and Baracchi, D. (2020). Exposure to a biopesticide interferes with sucrose responsiveness and learning in honey bees. Sci. Rep. 10, 19929 (2020).

Cepero, A., Ravoet, J., Gómez-Moracho, T., Bernal, J. L., Del Nozal, M. J., Bartolomé, C., Maside, X., Meana, A., González-Porto, A. V., de Graaf, D. C., Martín-Hernández, R. and Higes, M. (2014). Holistic screening of collapsing honey bee colonies in Spain: a case study. BMC Res. Notes. 7, 649 (2014).

Chapman, N. C., Colin, T., Cook, J., da Silva, C. R. B., Gloag, R., Hogendoorn, K., Howard, S. R., Remnant, E. J., Roberts, J. M. K., Tiertney, S. M., Wilson R. S. and Mikheyev, S. A. (2023). The final frontier: ecological and evolutionary dynamics of a global parasite invasion. Biol. Lett. 19:20220589.

Chen, P., Lu, Y.-H., Lin, Y.-H., Wu, C.-P., Tang, C.-K., Wei, S.-C. and Wu, Y.-L. (2021). Deformed wing virus infection affects the neurological function of Apis mellifera by altering extracellular adenosine signaling. Insect Biochem. Mol. Ecol., 139, 103674.

Chen, Y. R., Tzeng, D. T. W., Ting, C., Hsu, P. S., Wu, T. H., Zhong, S. and Yang, E. C. (2021). Missing Nurse Bees—Early Transcriptomic Switch From Nurse Bee to Forager Induced by Sublethal Imidacloprid. Front. Gen. 12. 10.3389/FGENE.2021.665927/FULL

Cox-Foster, D. L., Conlan, S., Holmes, E. C., Palacios, G., Evans, J. D., Moran, N. A., Quan, P. L., Briese, T., Hornig, M., Geiser, D. M., Martinson, V., VanEngelsdorp, D., Kalkstein, A. L., Drysdale, A., Hui, J., Zhai, J., Cui, L., Hutchison, S. K., Simons, J. F., … Lipkin, W. I. (2007). A metagenomic survey of microbes in honey bee colony collapse disorder. Science, 318(5848), 283–287.

de Miranda, J. R. and Genersch, E. (2010). Deformed wing virus. J. Invertebr. Pathol. 103(SUPPL. 1), S48–S61.

Decourtye, A., Lacassie, E. and Pham-Delégue, M. H. (2003). Learning performances of honeybees (Apis mellifera L) are differentially affected by imidacloprid according to the season. Pest Manag. Sci. 59(3), 269–278.

Deng, Y., Zhao, H., Yang, S., Zhang, L., Zhang, L. and Hou, C. (2020). Screening and Validation of Reference Genes for RT-qPCR Under Different Honey Bee Viral Infections and dsRNA Treatment. Front. Microbiol. 11, 1715.

Devaud, J.-M., Blunk, A., Podufall, J., Giurfa, M. and Grünewald, B. (2007). Using local anaesthetics to block neuronal activity and map specific learning tasks to the mushroom bodies of an insect brain. Eur. J. Neurosci. 26(11), 3193–3206.

Dolezal, A. G., Carrillo-Tripp, J., Judd, T. M., Allen Miller, W., Bonning, B. C. and Toth, A. L. (2019). Interacting stressors matter: diet quality and virus infection in honeybee health. R. Soc. Open Sci. 6(2).

Dupuis, J. P., Bazelot, M., Barbara, G. S., Paute, S., Gauthier, M. and Raymond-Delpech V. (2010). Homomeric RDL and Heteromeric RDL/LCCH3 GABA Receptors in the Honeybee Antennal Lobes: Two Candidates for Inhibitory Transmission in Olfactory Processing. J. Neurophysiol. 2010 103:1, 458-468.

Ganeshina, O. and Menzel, R. (2001), GABA-immunoreactive neurons in the mushroom bodies of the honeybee: An electron microscopic study. J. Comp. Neurol., 437: 335–349.

Gisder, S., Möckel, N., Eisenhardt, D. and Genersch, E. (2018). In vivo evolution of viral virulence: switching of deformed wing virus between hosts results in virulence changes and sequence shifts. Environ. Microbiol. 20(12), 4612–4628.

Giurfa, M. and Sandoz, J. C. (2012). Invertebrate learning and memory: Fifty years of olfactory conditioning of the proboscis extension response in honeybees. Learn. Mem. 19(2), 54–66.

Hellemans, J., Mortier, G., de Paepe, A., Speleman, F. and Vandesompele, J. (2007). qBase relative quantification framework and software for management and automated analysis of real-time quantitative PCR data. Genome Biol. 8(2), R19.

Homberg, U. (1984). Processing of antennal information in extrinsic mushroom body neurons of the bee brain. J. Comp. Physiol. A (1984) 154:825-836.

Huang, J., Wang, T., Qiu, Y., Hassanyar, A. K., Zhang, Z., Sun, Q., Ni, X., Yu, K., Guo, Y., Yang, C., Lü, Y., Nie, H., Lin, Y., Li, Z. and Su, S. (2023). Differential Brain Expression Patterns of microRNAs Related to Olfactory Performance in Honey Bees (Apis mellifera). Genes, 14(5).

Iqbal, J. and Mueller, U. (2007). Virus infection causes specific learning deficits in honeybee foragers. Proc. R. Soc. B. 274: 1517–1521.

Kanehisa, M. and Goto, S. (2000). KEGG: kyoto encyclopedia of genes and genomes. Nucleic Acids Res. 28(1), 27–30.

Lach, L., Kratz, M. and Baer, B. (2015). Parasitized honey bees are less likely to forage and carry less pollen. J. Inverteb. Pathol. 130, 64–71.

Liu, X. and Davis, R. L. (2008). The GABAergic anterior paired lateral neuron suppresses and is suppressed by olfactory learning. Nature Neurosci. 2008 12:1, 12(1), 53–59.

Liu, X., Krause, W. C. and Davis, R. L. (2007). GABAA Receptor RDL Inhibits Drosophila Olfactory Associative Learning. Neuron, 56(6), 1090–1102.

Mallon, E. B., Brockmann, A. and Schmid-Hempel, P. (2003). Immune response inhibits associative learning in insects. Proc. R. Soc. Lond. B. 270: 2471–2473.

Martin, S. J., Ball, B. V. and Carreck, N. L. (2015). The role of deformed wing virus in the initial collapse of varroa infested honey bee colonies in the UK. J. Apic. Res. 52(5): 251–258.

Martin, S. J. and Brettell, L. E. (2019). Deformed Wing Virus in Honeybees and Other Insects. Annu. Rev. Virol. Vol. 6:49–69.

Menzel, R. (2012). The honeybee as a model for understanding the basis of cognition. Nat. Rev. Neurosci. 13, 758–768 (2012).

Michels, B., Diegelmann, S., Tanimoto, H., Schwenkert, I., Buchner, E. and Gerber, B. (2005). A role for Synapsin in associative learning: The Drosophila larva as a study case. Learn. Mem. 12(3), 224–231.

Mota, T. and Giurfa, M. (2010). Multiple Reversal Olfactory Learning in Honeybees. Frontiers in Behav. Neurosci. 4(JUL).

Oldroyd, B. P. (2007). What’s Killing American Honey Bees? PLoS Biol. 5(6), e168.

Pizzorno, M. C., Field, K., Kobokovich, A. L., Martin, P. L., Gupta, R. A., Mammone, R., Rovnyak, D. and Capaldi, E. A. (2021). Transcriptomic Responses of the Honey Bee Brain to Infection with Deformed Wing Virus. Viruses 2021, 13(2), 287.

R Core Team. (2017). R: A language and environment for statistical computing. R Foundation for Statistical Computing.

Raccuglia, D. and Mueller, U. (2013). Focal uncaging of GABA reveals a temporally defined role for GABAergic inhibition during appetitive associative olfactory conditioning in honeybees. Learn. Mem. 20(8), 410–416.

Sadanandappa, M. K., Redondo, B. B., Michels, B., Rodrigues, V., Gerber, B., VijayRaghavan, K., Buchner, E. and Ramaswami, M. (2013). Synapsin function in GABA-ergic interneurons is required for short-term olfactory habituation. J. Neurosci. 33(42), 16576–16585.

Scheiner, R., Barnert, M. and Erber, J. (2003). Variation in water and sucrose responsiveness during the foraging season affects proboscis extension learning in honey bees. Apidologie, 34(1), 67–72.

Shah, K. S., Evans, E. C. and Pizzorno, M. C. (2009). Localization of deformed wing virus (DWV) in the brains of the honeybee, Apis mellifera Linnaeus. Virol. J. 6(1), 1–7.

Siviter, H., Koricheva, J., Brown, M. J. F. and Leadbeater, E. (2018). Quantifying the impact of pesticides on learning and memory in bees. J. Appl. Ecol. 2018; 55: 2812–2821.

Tang, C. K., Lin, Y. H., Jiang, J. A., Lu, Y. H., Tsai, C. H., Lin, Y. C., Chen, Y. R., Wu, C. P. and Wu, Y. L. (2021). Real-time monitoring of deformed wing virus-infected bee foraging behavior following histone deacetylase inhibitor treatment. iScience, 24(10), 103056.

Tavares, D. A., Roat, T. C., Silva-Zacarin, E. C. M., Nocelli, R. C. F. and Malaspina, O. (2019). Exposure to thiamethoxam during the larval phase affects synapsin levels in the brain of the honey bee. Ecotoxicol. Environ. Saf. 169, 523–528.

VanderWeele, T. J. and Mathur, M. B. (2019). SOME DESIRABLE PROPERTIES OF THE BONFERRONI CORRECTION: IS THE BONFERRONI CORRECTION REALLY SO BAD? Am. J. Epidemiol. 188(3), 617–618.

Vandesompele, J., de Preter, K., Pattyn, F., Poppe, B., van Roy, N., de Paepe, A. and Speleman, F. (2002). Accurate normalization of real-time quantitative RT-PCR data by geometric averaging of multiple internal control genes. Genome Biol. 3(7), research0034.1.

Veiner, M., Morimoto, J., Leadbeater, E. and Manfredini, F. (2022). Machine learning models identify gene predictors of waggle dance behaviour in honeybees. Mol. Ecol. Resour. 22(6), 2248–2261.

Wells, T., Wolf, S., Nicholls, E., Groll, H., Lim, K. S., Clark, S. J., Swain, J., Osborne, J. L. and Haughton, A. J. (2016). Flight performance of actively foraging honey bees is reduced by a common pathogen. Environ. Microbiol. Rep. 8(5), 728–737.

Woodford, L., Christie, C. R., Campbell, E. M., Budge, G. E., Bowman, A. S. and Evans, D. J. (2022). Quantitative and Qualitative Changes in the Deformed Wing Virus Population in Honey Bees Associated with the Introduction or Removal of Varroa destructor. Viruses, 14(8).

Zhang, W., Wang, L., Zhao, Y., Wang, Y., Chen, C., Hu, Y., Zhu, Y., Sun, H., Cheng, Y., Sun, Q., Zhang, J. and Chen, D. (2022). Single-cell transcriptomic analysis of honeybee brains identifies vitellogenin as caste differentiation-related factor. iScience, 25(7).

